# Spatial analysis of the cure rate for tuberculosis in primary health care in the municipality of Rio de Janeiro between 2012 and 2014

**DOI:** 10.1101/340752

**Authors:** José Carlos Prado Junior, Roberto de Andrade Medronho

## Abstract

**Background:** Tuberculosis (TB) has a high disease burden and the World Health Organization (WHO) states it is a global emergency. TB is the most important cause of death from infectious disease in adults. It is directly related to access to health services and socioeconomic factors. Primary health care (PHC) provides greater linkage of people to health services and greater medication adherence in some chronic diseases. It also provides supervised treatment and more effective search for contactants. The PHC Reform started in 2009 in Rio de Janeiro, increasing coverage from 7% to 46.16% in 2015.

**Methodology/Principal findings:** This paper aims to evaluate the spatial distribution of new TB cases closed with a cure outcome in dwellers of Rio de Janeiro in the period 2012-2014, according to PHC coverage, controlling socioeconomic, demographic and epidemiological factors. Variables were obtained from the Notifiable Diseases Information System for Tuberculosis (SINAN-TB) and the socioeconomic variables from the 2010 national census at census tract level. The socioeconomic variables were selected from multivariate analysis using main factors analysis technique. The generalized additive model (GAM) was used for the spatial analysis. Association was found between TB cure and variables education, alcoholism, contacts search, HIV serology and elderly. People with family health coverage between 35 and 41 months were 1.64 more likely of cure when compared to people without coverage (95% CI 1.07-2.51). Spatial analysis identified areas with less probability of cure for tuberculosis in the municipality of Rio de Janeiro.

**Author Summary:** Tuberculosis is associated to social and demographic conditions. Lack of access to healthcare contributes to delay in diagnosis and in the beginning of the treatment. Primary health care improve access and adherence to treatment. This study can be useful as a public health policy, since it is possible to prioritize the region in the map to improve TB cure. We found association between tuberculosis cure and the duration of implantation of the primary health care teams. This finding corroborates the importance of treating tuberculosis in this level of care. The spatial analysis of cases of tuberculosis cure showed a significant spatial association with the cure of tuberculosis. The results of this study can contribute reinforcing the policy makers for developing primary health care to improve the access to health services and to reach better TB cure rates. Spatial analysis may be an useful tool for identifying the areas where to prioritize efforts for reaching better results.

## Introduction

Tuberculosis (TB) is an infectious disease of great magnitude and importance in the world. It was declared a worldwide emergency disease by the World Health Organization (WHO) in 1993 and is the largest cause of death by infectious diseases in adults [1].

Brazil is one of WHO 22 priority TB control countries and ranked 16^th^ in 2014. These countries together account for 80% of global cases. TB prevalence in Brazil in 2014 was 33.5 / 100,000 inhabitants and mortality rate (MR) was 2.3 deaths / 100,000 inhabitants. Despite this position, an average annual incidence reduction of 2.3% and 0.5 deaths / 100,000 inhabitants has been recorded between 2005 and 2014 [1].

In 2014, the incidence rate in the municipality of Rio de Janeiro was 66.8 / 100,000 inhabitants. The cure rate of smear-positive pulmonary tuberculosis in the city was 69.2%, below that recommended by the WHO, which is 85% [2].

The disease has a direct relationship with the misery and social exclusion, therefore the most socially vulnerable people have a greater probability of developing and proliferating this disease.

Socioeconomic factors hamper people’s access to health services [3] which in turn contribute to delayed TB diagnosis and treatment, increasing the possibility of abandoning treatment, one of the main obstacles to control this disease [3].

PHC plays a key role in curing tuberculosis because it is the first level of access of a health system, with some basic principles of organization such as longitudinality, comprehensive care and coordination of care within the system itself [4, 5]. Therefore, PHC allows greater access and adherence to TB treatment, as well as reaching the most vulnerable populations [4].

The countries that adhered to PHC as a health system organizer showed evidence of improvement in care indicators as well as lower health investments when compared to those without structured PHC [4,6,7,8].

An association between PHC expanded coverage and reduced treatment abandonment rate and lower mortality due to tuberculosis is described [9].

The Municipality of Rio de Janeiro (MRJ) had ESF coverage of approximately 7% up to 2008 [10] and only 3.55% coverage of full teams [11, 12]. As of 2009, the PHC reform began with expanded coverage to approximately 40% in 2012 and 46.16% in 2015, considering full teams [6, 10].

Currently, there are two PHC care models: (1) Family Health, with a more comprehensive supply of health services based on PHC principles (access, longitudinality, care coordination and comprehensiveness) [4] with general practitioners and nurses with territorially-defined health responsibility; and (2) Traditional PHC, with medical care based on gender and age (pediatrics, gynecology, medical clinic) [6].

Considering that TB is a disease strongly related to socioeconomic factors [13,14,15,16,17,18,19], as well as to access to health services[3, 9], it is necessary to evaluate the relationship of expanded PHC coverage in the cure of TB in the municipality.

This paper aims to evaluate the spatial distribution of new cases of tuberculosis closed with a cure outcome in residents of Rio de Janeiro in the period 2012-2014, by PHC coverage, controlling socioeconomic, demographic and epidemiological factors.

## Methods

This cross-sectional study was carried out in Rio de Janeiro, correlating TB cure with family health coverage and socioeconomic, demographic and epidemiological variables, and the unit of analysis is made of individuals from newly reported cases of tuberculosis in the period 2012-2014. The municipality is located in the southeast of the country at geographic coordinates 22°44′45.59″ to 23°04′58.34″ south latitude and 43°05′48.89″ to 43°47′43.79″ west longitude; it has an area of 1,199.82 km^2^, a population density of 5,265.82 inhabitants / km^2^ and an exclusively urban population estimated at 6,476,631 inhabitants in 2015 [20]. It is divided into 160 neighborhoods, 33 administrative regions and 10 health districts (planning areas). Rio shows great economic and social contrasts, with approximately 22% of the population residing in subnormal clusters [21].

The study population is comprised of new cases of tuberculosis residing in the MRJ notified in the period 2012-2014. All cases were eligible to be part of the study in order to minimize selection bias.

The geo-referencing technique was used from the residence address to identify family’s health coverage, assigning a geographical position for each record (latitude and longitude). Thus, it was possible to identify through spatial consultation the family health’s coverage polygons to which the point was inserted. The coverage maps were developed by the Municipal Health Secretariat of Rio de Janeiro and correspond to the existing coverage in 2012, 2013 and 2014, with reference to the month of December. Georeferencing of addresses was done using the “Geocode” tool made available by Google Maps through a free Application Programming Interface (API).

This process utilized the Google’s streets and location base. The accuracy of georeferencing can be evaluated from a score ranging from 0 to 10, (0-not found, 1-country level, 2-state, 3-subregion, 4-city, 5-Zip Code, 6-streets, 7-intersection between streets, 8-address, 9-name of the building or trade, 10-maximum precision). Addresses with a score lower than “5” were considered as losses. Records with scores between “8” and “10” were considered with acceptable accuracy. The remaining records were manually reviewed. Of the 14,384 georeferenced records, the precise geographic coordinates of 3,484 records (24.22%) could not be determined, characterizing losses of the georeferenced sample. The total number of georeferenced records was 10,900 new closed cases.

The outcome variable considered in this study was the “closed with a cure outcome” (yes / no) obtained from the SINAN-TB and the exposure variable was “family health coverage”, expressed by the “time of implantation of the teams until TB diagnosis (in months)”.

The selection of variables was based on the theoretical TB cure model (Fig 1) based on the realms “environment”, “individual factors”, “access to health service” and “social status”. The model includes demographic, social, epidemiological, access and use of health services variables.

**Fig 1.**
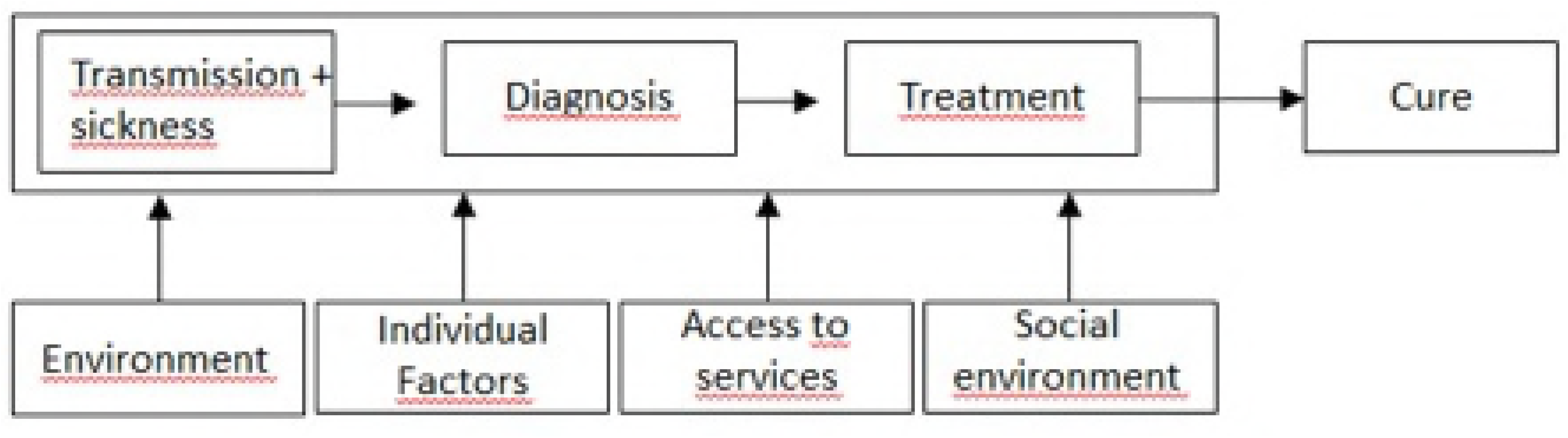
Model of analysis for the cure of tuberculosis ^a^. ^a^ To cure tuberculosis, an individual with the disease must have been diagnosed and treatment performed. The environment, individual factors, access to health services and social condition (agglomeration, income, education, household conditions and demographic factors) of the individual influence the transmission, diagnosis and treatment of tuberculosis. Proper treatment results in the cure of tuberculosis.

The variable “family health teams implantation time” reflects access to health service and was obtained from data from the National Registry System of Health Establishments of the Ministry of Health (SCNES). For the purpose of classifying family health coverage, all records located in polygons of areas with family health coverage whose diagnosis was given after 3 months of implantation of the team were considered with coverage and all other cases without family health coverage. The 3-month period was used as cut-off point because it is the minimum reasonable time for a new family health team to carry out all registration of people residing in its health territory and produce an initial health diagnosis.

The variables selected from the SINAN-TB were: age; gender; race / color (white / nonwhite); schooling; “HIV coinfection” (yes / no); “alcohol abuse history” (yes / no); “Contact search” (yes / no); “serology for HIV” (positive, negative, not performed); “supervised treatment” (yes / no).

Considering that there is a large number of socioeconomic variables from the 2010 demographic census (IBGE, 2011)^22^, a multivariate analysis was performed using the main components analysis technique. Thus, the TB cure-related realms (“agglomeration”, “household conditions”, “demographic”, “schooling”, “income”) were established from the theoretical model related to the “social condition” (Fig 1) and for each realm, the variables that, to a lesser extent, represent the other variables of that realm were selected.

The socioeconomic and demographic variables resulting from this analysis were: “average monthly income of the person in charge (R$)”; “average number of residents per household”; “population density in the census sector”; “Density of dwellers/rooms”; “proportion of permanent private households with bathrooms for the exclusive use of residents or water closet and sanitary sewage via general sewer or rainwater network”; “Proportion of permanent households with electricity”; “average number of toilets per permanent private residence”; “aging rate”.

The variables obtained from the Demographic Census represent the averages and proportions of each census tract. These values were repeated for each individual resident in the same census tract, since these variables in this level of aggregation show high homogeneity. The other variables were analyzed from the individual level.

Data descriptive analysis was performed using the R software (R Development Core Team, 2016)^23,24^ through bivariate logistic regression (gross analysis).

During TB spatial exploratory analysis, we observed the distribution of points and identification of possible clusters through the point density estimation technique, defined as Kernel density estimation, which consists of generating a point density surface within a region of influence, weighted by the distance of each from the location of interest, for the visual identification of “hot areas” on the map. Bandwidths from 500m to 3,000m were tested, with 250m increments. A matrix of 500×500 points and a radius of 2,500 m was used because it was considered the most appropriate for highlighting strategic areas. We generated maps with estimates of TB incidence rate through the kernel ratio between reported cases of TB and the kernel of the population.

Spatial analysis was based on the generalized additive model (GAM), which can be considered an extension of generalized linear models, with the inclusion of a non-parametric element by smoothing functions. The model has the great advantage of being more flexible and relatively simple to interpret [25]. The Thin Plate Regression Splines smoothing technique was used.

The construction of the generalized additive models was performed through manual selection, based on the essential factors of the theoretical framework (Fig 1), not only considering the statistically significant variables in the bivariate analysis, but also those of epidemiological importance.

We did not only consider the p-value of each association, but the importance previously described of each variable and the impact on the explanatory power of the model. Only those variables with a clear negative impact under the explanatory power of the model, observed through deviance, were removed.

The software for constructing family health coverage maps and spatial data queries was ArcGIS, version 10.2.2 [26] in Latlong/WGS84 projection, available in the shapefile extension.

R software [23] was used with RStudio software [24], version 0.99.893 for the exploratory analysis, multivariate analysis, kernel estimation and spatial analysis. During the analyses, the following additional packages of R were used: car [27], mgcv [28], descr [29], sp [30], sdep [31], maptools [32], splancs [33], fields [34], RColorBrewer [35], ggplot2 [36].

This work was funded by the authors. The Ethics Committee of the Municipal Health Secretariat of Rio de Janeiro approved the study, under opinion N° 1.389.137 and CAAE protocol 52493216.5.0000.5279, in compliance with the recommendations contained in Resolution 466/12 of the National Health Council. All data analyzed were anonymized.

## Results

During the years 2012 to 2014, 16,363 new TB cases of MRJ residents were identified. Of these, 905 cases were excluded from the study due to lack of closure in addition to 161 cases of street population and other 913 records of institutionalized persons, since these individuals cannot be classified in terms of family health coverage, leaving out 14,384 records for the study.

The closing percentage for this period was 94.47%. The cure rate was 71.57% (11,063 new cases) of the total cases closed. The mean incidence rate was 84.91 cases / 100,000 inhabitants. In the same period, there were 725 cases of death by TB resulting in a mean specific mortality rate of 3.76 cases per 100,000 inhabitants, and the case fatality rate was 4.69% among new cases closed. Median age was 38 years.

The estimation of the incidence rate from the Kernel ratio (Fig 2) shows that the “hot areas” are mainly concentrated in the southern zone (Rocinha), followed by the northern zone (Complexo do Alemão, Acari, Pavuna) and the western zone (Senador Camará, Realengo).

**Fig 2.**
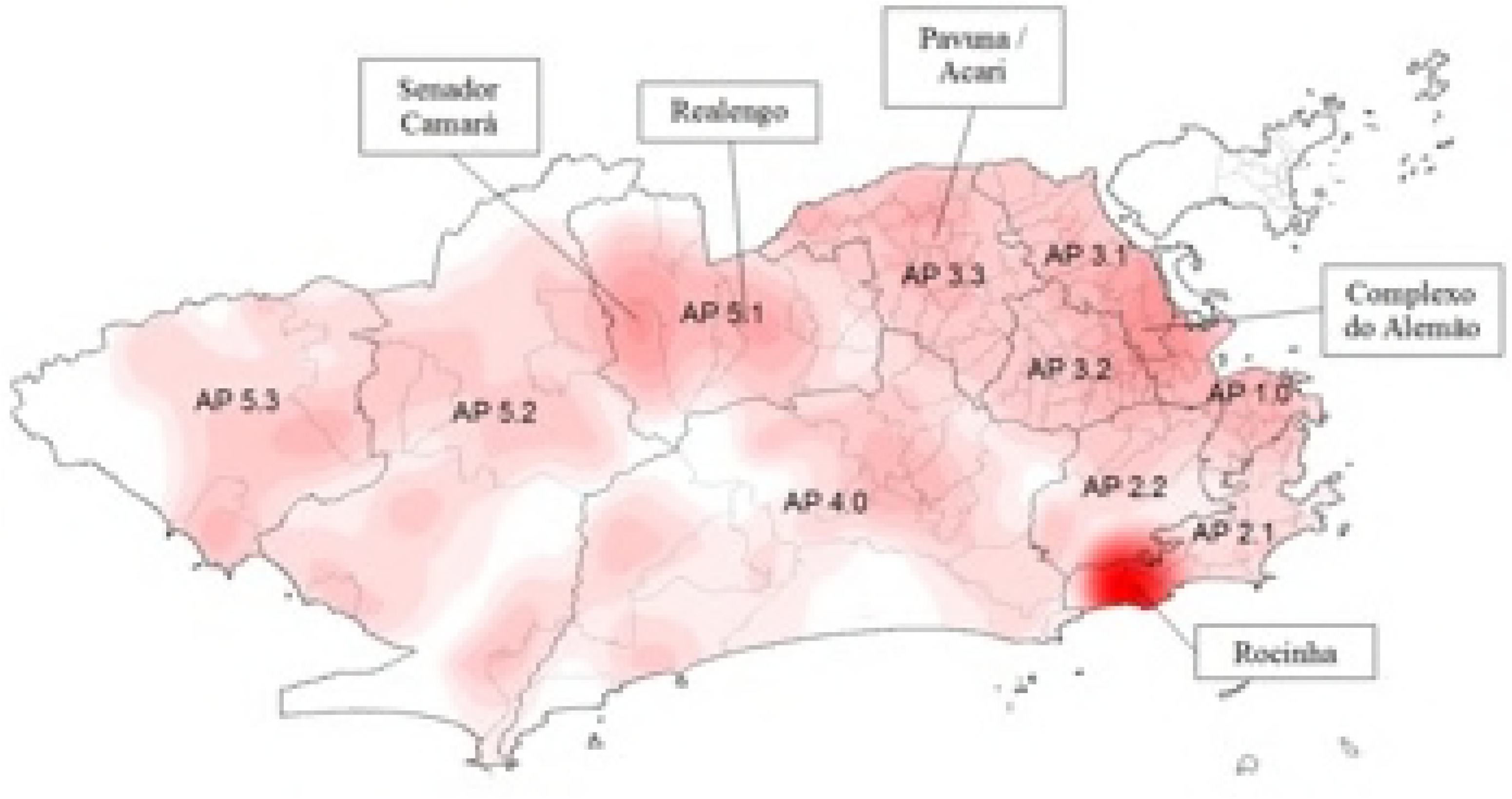
Kernel ratio among new cases of tuberculosis and population. Municipality of Rio de Janeiro (RJ), Brazil, from 2012 to 2014. Source: Cartographic data: IPP, Epidemiological data: SINAN-TB. Projection GCS. Datum WGS84

Of the new TB cases between 2012 and 2014 that were part of the study (n = 14,384), most cases occurred in males (n = 8,919; 62.0%), of non-white race / color = 7,839, 54.5%) and illiterates (n = 4,493, 31.2%).

An association with several variables was found (Table 1) from the crude bivariate analysis of TB cure with socioeconomic and demographic variables. In relation to age, there is a lower probability of cure in the age range > 25 to 50 years (OR 0.86, 95% CI 0.78-0.94) compared to the reference category (0 to 25 years). Among individuals aged > 80 years, the probability of cure decreases even further (OR 0.44, 95% CI 0.32-0.59). Women were 1.51 times (95% CI 1.39-1.63) more likely to be cured than men. Non-white people were less likely to be cured (OR 0.68, 95% CI 0.62-0.73) than white people. The higher the level of schooling, the greater the probability of cure compared to illiterate people. The higher the average monthly income of the person responsible, the greater the probability of cure. On the other hand, the higher the average number of residents in permanent private households, the lower the probability of cure. The greater the percentage of permanent private households with electricity and the average number of bathrooms per household, the greater the probability of a cure. The higher the aging rate of the census tract of the individual, the greater the probability of cure, evidencing an apparent paradox with findings in the age groups measured at the individual level.

**Table 1.**
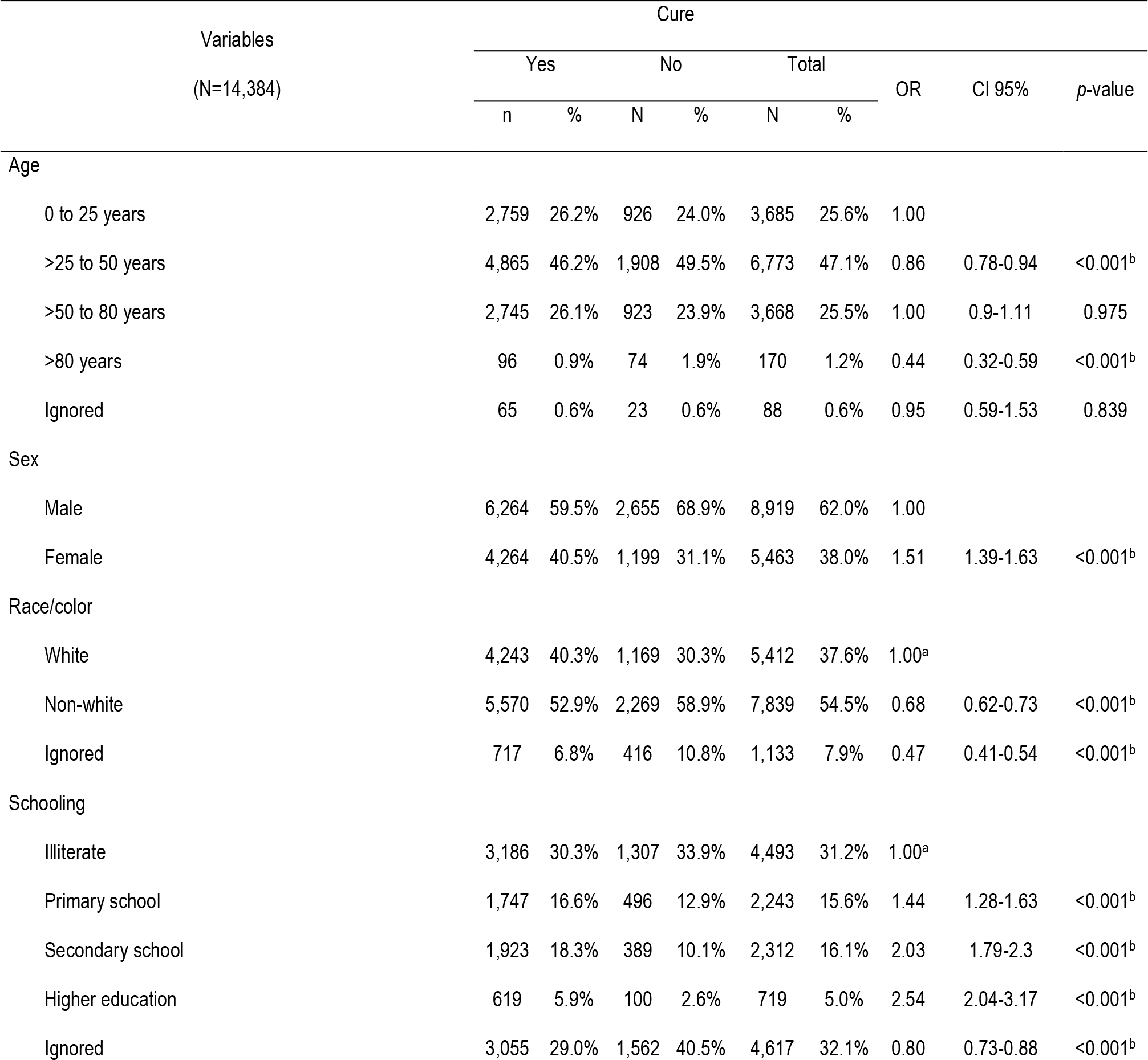
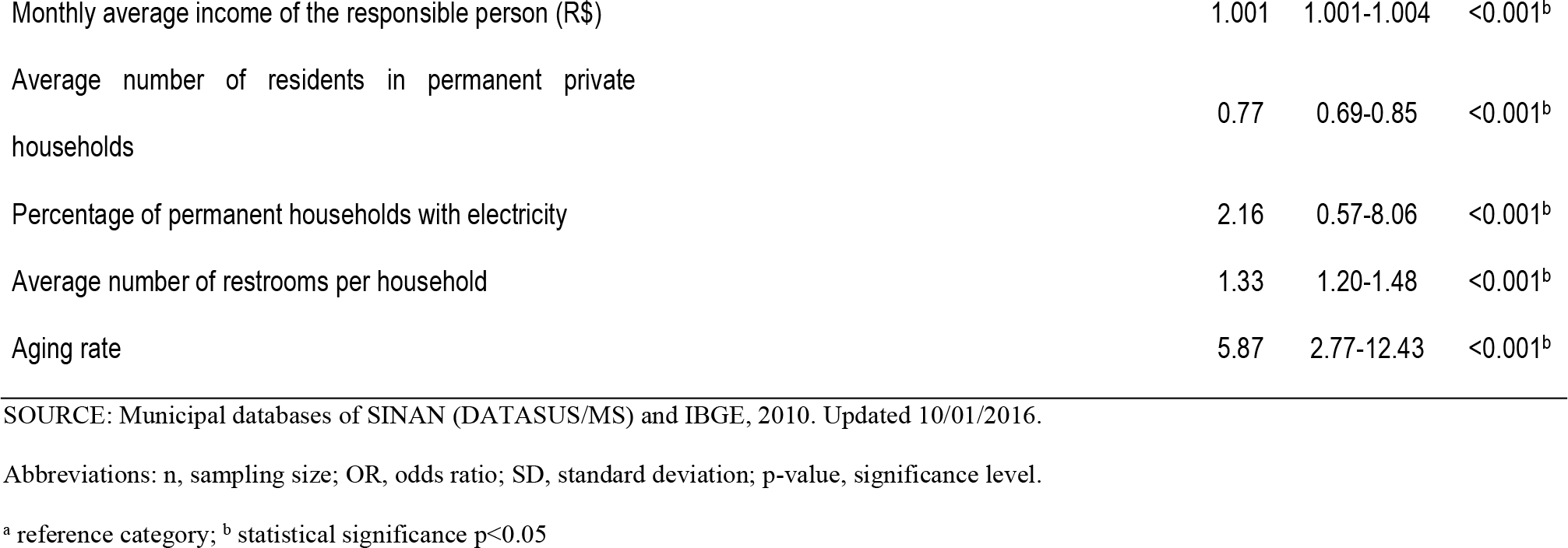
Gross cure analysis of new cases of tuberculosis with socioeconomic variables (N = 14,384). Municipality of Rio de Janeiro (RJ), Brazil, 2012-2014.

| Variables<br>(N=14,384) |  | Cure |  |  |  |  |  | OR | CI 95% | p-value |
| --- | --- | --- | --- | --- | --- | --- | --- | --- | --- | --- |
|  |  | Yes |  | No |  | Total |  |  |  |  |
|  |  | n | % | N | % | N | % |  |  |  |
| Age |  |  |  |  |  |  |  |  |  |  |
| 0 to 25 years |  | 2,759 | 26.2% | 926 | 24.0% | 3,685 | 25.6% | 1.00 |  |  |
| >25 to 50 years |  | 4,865 | 46.2% | 1,908 | 49.5% | 6,773 | 47.1% | 0.86 | 0.78-0.94 | <0.001<sup>b</sup> |
| >50 to 80 years |  | 2,745 | 26.1% | 923 | 23.9% | 3,668 | 25.5% | 1.00 | 0.9-1.11 | 0.975 |
| >80 years |  | 96 | 0.9% | 74 | 1.9% | 170 | 1.2% | 0.44 | 0.32-0.59 | <0.001<sup>b</sup> |
| Ignored |  | 65 | 0.6% | 23 | 0.6% | 88 | 0.6% | 0.95 | 0.59-1.53 | 0.839 |
| Sex |  |  |  |  |  |  |  |  |  |  |
| Male |  | 6,264 | 59.5% | 2,655 | 68.9% | 8,919 | 62.0% | 1.00 |  |  |
| Female |  | 4,264 | 40.5% | 1,199 | 31.1% | 5,463 | 38.0% | 1.51 | 1.39-1.63 | <0.001<sup>b</sup> |
| Race/color |  |  |  |  |  |  |  |  |  |  |
| White |  | 4,243 | 40.3% | 1,169 | 30.3% | 5,412 | 37.6% | 1.00<sup>a</sup> |  |  |
| Non-white |  | 5,570 | 52.9% | 2,269 | 58.9% | 7,839 | 54.5% | 0.68 | 0.62-0.73 | <0.001<sup>b</sup> |
| Ignored |  | 717 | 6.8% | 416 | 10.8% | 1,133 | 7.9% | 0.47 | 0.41-0.54 | <0.001<sup>b</sup> |
| Schooling |  |  |  |  |  |  |  |  |  |  |
| Illiterate |  | 3,186 | 30.3% | 1,307 | 33.9% | 4,493 | 31.2% | 1.00<sup>a</sup> |  |  |
| Primary school |  | 1,747 | 16.6% | 496 | 12.9% | 2,243 | 15.6% | 1.44 | 1.28-1.63 | <0.001<sup>b</sup> |
| Secondary school |  | 1,923 | 18.3% | 389 | 10.1% | 2,312 | 16.1% | 2.03 | 1.79-2.3 | <0.001<sup>b</sup> |
| Higher education |  | 619 | 5.9% | 100 | 2.6% | 719 | 5.0% | 2.54 | 2.04-3.17 | <0.001<sup>b</sup> |
| Ignored |  | 3,055 | 29.0% | 1,562 | 40.5% | 4,617 | 32.1% | 0.80 | 0.73-0.88 | <0.001<sup>b</sup> |

|  |  |  |  |
| --- | --- | --- | --- |
| Monthly average income of the responsible person (R\$) | 1.001 | 1.001-1.004 | <0.001<sup>b</sup> |
| Average number of residents in permanent private households | 0.77 | 0.69-0.85 | <0.001<sup>b</sup> |
| Percentage of permanent households with electricity | 2.16 | 0.57-8.06 | <0.001<sup>b</sup> |
| Average number of restrooms per household | 1.33 | 1.20-1.48 | <0.001<sup>b</sup> |
| Aging rate | 5.87 | 2.77-12.43 | <0.001<sup>b</sup> |

SOURCE: Municipal databases of SINAN (DATASUS/MS) and IBGE, 2010. Updated 10/01/2016.
Abbreviations: n, sampling size; OR, odds ratio; SD, standard deviation; p-value, significance level.
<sup>a</sup> reference category; <sup>b</sup> statistical significance p<0.05

An association between cure of new tuberculosis cases and some epidemiological variables was found (Table 2). People without AIDS coinfection were 3.46 (95% CI: 3.07-3.91) times more likely of being cured when compared to people with coinfection. Similarly, people with negative HIV serology were 3.67 (95% CI 3.28-4.12) times more likely of being cured compared to people with positive serology. On the other hand, people with unsupervised treatment were less likely to be cured (OR 0.65, 95% CI 0.60-0.70) compared with those whose treatment was supervised, and people whose cases had no search for contactants were also less likely to be cured (OR 0.36, 95% CI 0.33-0.39) compared to those who did. In relation to the Family Health coverage, a slightly higher probability of cure is perceived the higher the coverage, but without statistical significance.

**Table 2.** Gross cure analysis of new cases of tuberculosis with epidemiological and healthcare variables (N = 14,384). Municipality of Rio de Janeiro (RJ), Brazil, 2012-2014.

| Variables<br>(N=14,384) | Cure |  |  |  |  |  | OR | CI 95% | p-value |
| --- | --- | --- | --- | --- | --- | --- | --- | --- | --- |
|  | Yes |  | No |  | Total |  |  |  |  |
|  | n | % | N | % | N | % |  |  |  |
| AIDS Coinfections |  |  |  |  |  |  |  |  |  |
| Yes | 606 | 8.0% | 643 | 23.1% | 1,249 | 12.1% | 1.00 |  |  |
| No | 6,977 | 92.0% | 2,139 | 76.9% | 9,116 | 87.9% | 3.46 | 3.07-3.91 | <0.001<sup>b</sup> |
| HIV test performed |  |  |  |  |  |  |  |  |  |
| Positive | 744 | 7.1% | 707 | 18.3% | 1,451 | 10.1% | 1.00 |  |  |
| Negative | 7,573 | 71.9% | 1,960 | 50.9% | 9,533 | 66.3% | 3.67 | 3.28-4.12 | <0.001<sup>b</sup> |
| Not performed | 2,213 | 21.0% | 1,187 | 30.8% | 3,400 | 23.6% | 1.77 | 1.56-2.01 | <0.001<sup>b</sup> |
| Supervised treatment |  |  |  |  |  |  |  |  |  |
| Yes | 6.181 | 65.2% | 1,895 | 54.9% | 8,076 | 62.4% | 1.00 |  |  |
| No | 3.300 | 34.8% | 1,556 | 45.1% | 4,856 | 37.6% | 0.65 | 0.6-0.7 | <0.001<sup>b</sup> |
| Search for contactants |  |  |  |  |  |  |  |  |  |
| Yes | 8.572 | 81.4% | 2,359 | 61.2% | 10,931 | 76.0% | 1.00 |  |  |
| No | 1.958 | 18.6% | 1,495 | 38.8% | 3,453 | 24.0% | 0.36 | 0.33-0.39 | <0.001<sup>b</sup> |
| Alcohol abuse |  |  |  |  |  |  |  |  |  |
| Yes | 769 | 8.7% | 524 | 17.2% | 1,293 | 10.9% | 1.00 |  |  |
| No | 8.066 | 91.3% | 2,517 | 82.8% | 10,583 | 89.1% | 2.18 | 1.94-2.46 | <0.001<sup>b</sup> |
| FH coverage |  |  |  |  |  |  |  |  |  |
| 4 to 18 months | 1.211 | 26.8% | 417 | 25.8% | 1,628 | 26.6% | 1.00 |  |  |
| >18 to 35 months | 1.766 | 39.1% | 680 | 42.0% | 2,446 | 39.9% | 0.89 | 0.78-1.03 | 0.115 |
| >35 to 41 months | 418 | 9.3% | 124 | 7.7% | 542 | 8.8% | 1.16 | 0.92-1.46 | 0.207 |
| >41 months | 1.118 | 24.8% | 397 | 24.5% | 1,515 | 24.7% | 0.97 | 0.83-1.14 | 0.717 |

SOURCE: Municipal databases of SINAN (DATASUS/MS) and IBGE, 2010. Updated 10/01/2016.
Abbreviations: n, sampling size; OR, odds ratio; SD, standard deviation; p-value, significance level.
<sup>a</sup> reference category; <sup>b</sup> statistical significance p<0.05

The spatial Generalized Additive Model (GAM) enabled the building of a TB cure probability map. Thus, it was assumed that cases were all newly reported cases of tuberculosis in MRJ in the period from 2012 to 2014 that were not closed due to cure outcome (non-cure), and controls those records closed due to cure outcome (Fig 3).

**Fig 3.**
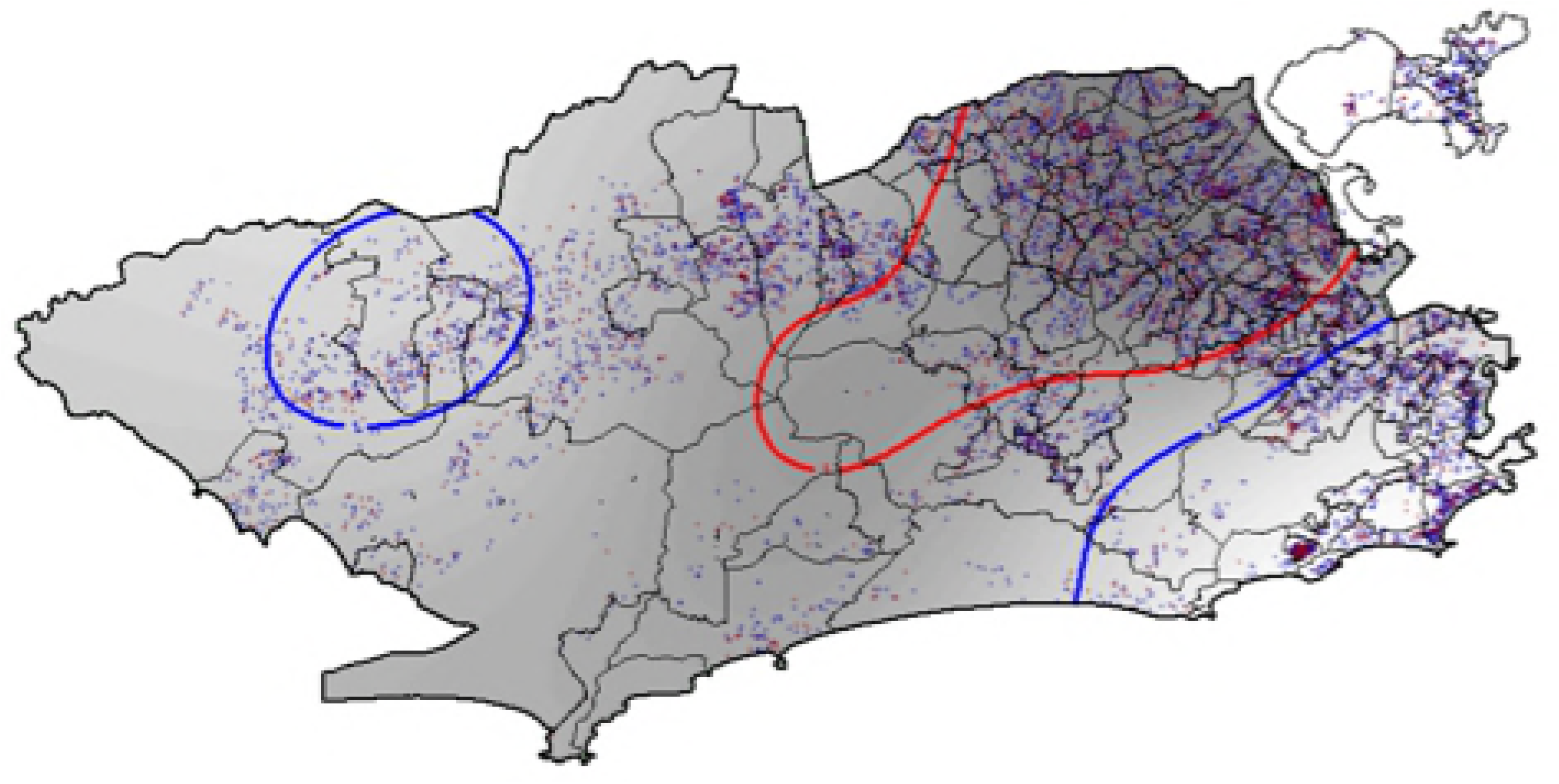
Map of probability of cure of tuberculosis using the Generalized Additive Model – GAM for cases (not cured) and controls (cure). Rio de Janeiro, 2012 to 2014. Source: Cartographic data: IPP, Epidemiological data: SINAN-TB. Projection GCS. Datum WGS84

It is possible to observe that people residing within the blue contours in the west and sou
th zones (right and left ends of the map) represent areas with an OR significantly above 1, that is, people residing in those areas tend to have more tuberculosis cure, whereas those living in the north, surrounded by the red contour (central contour of the map) tend to have less probability of cure.

The final model of the spatial analysis of new TB cases in MRJ in the years 2012 to 2014 for TB cure and the socioeconomic, demographic and epidemiological variables using the generalized additive model is presented in Table 3.

**Table 3.**
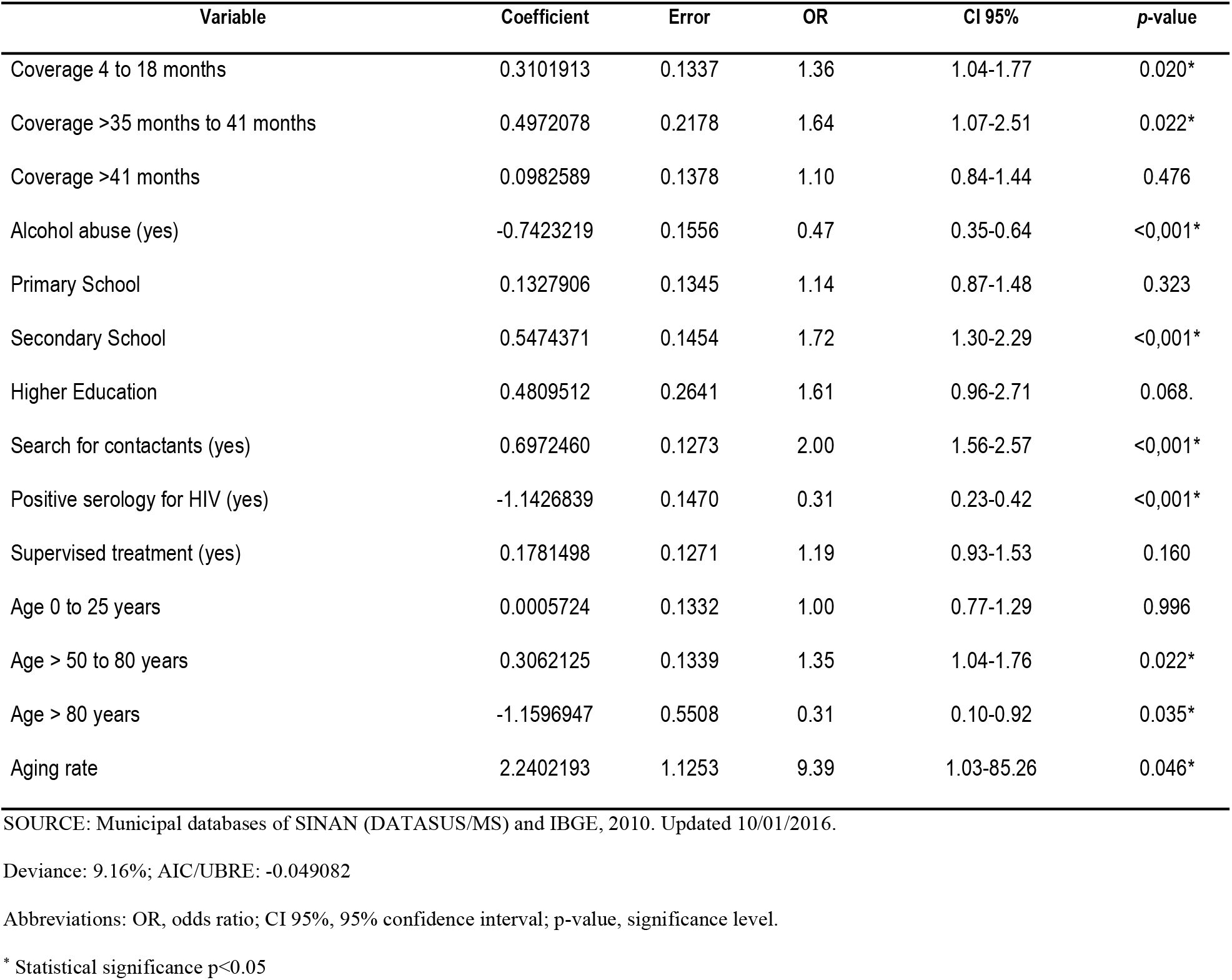
Final model of spatial analysis of new cases of tuberculosis with socioeconomic, demographic and epidemiological variables using the generalized additive model - GAM (N = 14,384). Municipality of Rio de Janeiro (RJ), Brazil, 2012-2014.

The family health coverage variable, measured here as time lapse of teams implantation at the time of diagnosis, was significant for the categories “4 to 18 months” and “> 35 months to 41 months”. People covered by family health tend to cure more for tuberculosis the longer the team’s time of implantation: OR 1.36 (95% CI 1.04-1.77) and OR 1.64 (95% CI 1.07-2, 51), respectively. People with a history of alcohol abuse had almost half the probability of cure (OR 0.47, 95% CI 0.35-0.64) when compared to people with no history of alcohol abuse. There was a positive trend in the association between the educational level and the probability of TB cure, especially in the case of secondary school students, with 1.72 times the probability of cure (95% CI 1.30-2.29) compared to illiterate people. People where the search for contactants occurred were 2.00 (95% CI 1.56-2.57) times more likely of achieving cure. People who had positive HIV serology were 0.31 (95% CI 0. 23-0.42) times less likely of achieving cure. On the other hand, people residing in census tracts with higher aging rates were 9.39 (95% CI 1.03-85.26) times more likely of achieving cure. The figure below illustrates the same final spatial model through a chart with the odds ratio of the variables.

**Fig 4.**
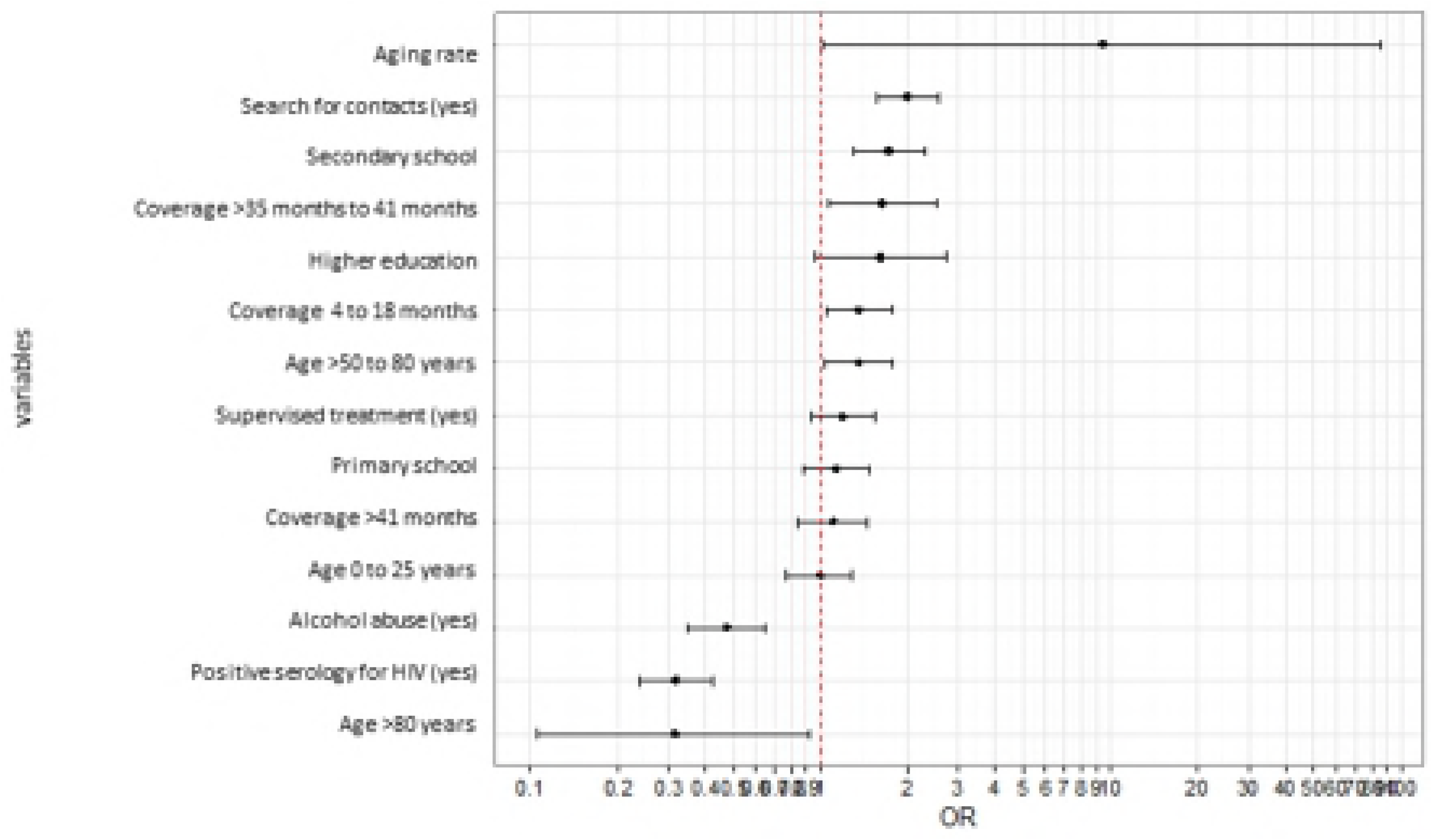
Final spatial analysis model using the generalized additive model - GAM for non-cure of tuberculosis. Rio de Janeiro, 2012 to 2014. The final spatial model map can be visualized in Fig 5. It is the smoothed spatial component, adjusted for the other socioeconomic, demographic and epidemiological variables of the final model. Significant spatial association was found (p = 0.0219). The areas surrounded by green dotted lines had positive spatial correlation for cure, whereas red dotted lines had inverse spatial correlation with cure.

**Fig 5.**
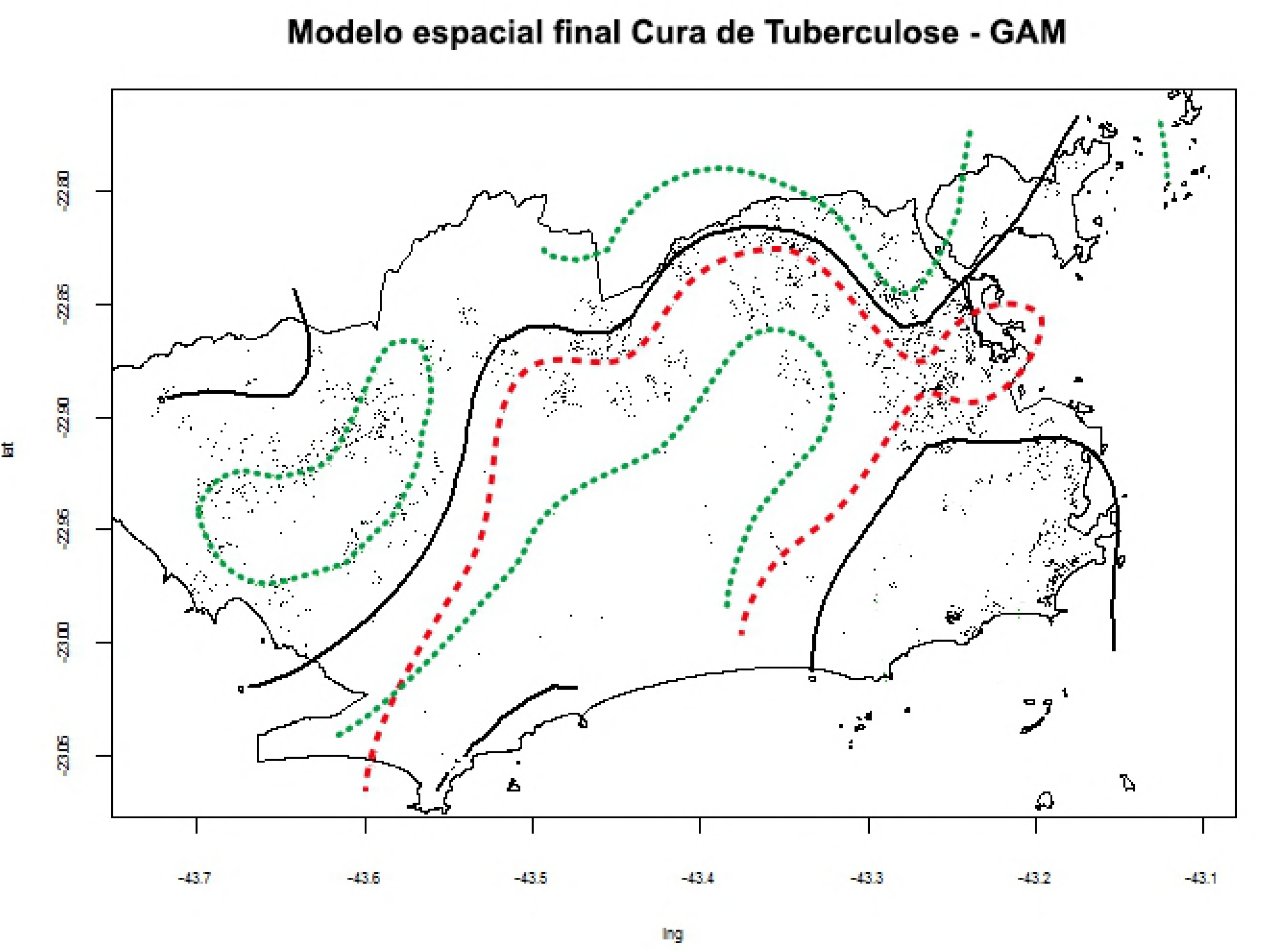
Final spatial analysis model using the generalized additive model - GAM for tuberculosis cure - smoothed spatial component adjusted for the other covariates. Rio de Janeiro, 2012 to 2014.

## DISCUSSION

In Brazil, there are few studies on the spatial distribution of endemic diseases, such as TB, in urban areas [15]. The information about the spatial and temporal spread of these diseases allows to understand the occurrence of these events in the territory. In addition, the description and visualization of the spatial distribution of the event facilitate the identification of its association with local characteristics, such as socioeconomic conditions.

The cure rate of 71.57% was lower than the value found in other studies conducted in other regions of the country, such as Silva et al (2014)[17], who found 90.9% of the cases closed due to cure in Maranhão. However, it is better than in Crato (CE), where Pinto et al (2015)[18] found a cure rate of 47.2%.

Spatial analysis demonstrated a significant spatial association with TB cure. In addition, the TB cure probability map shows that the southern and western zones were more likely to be cured from tuberculosis while residents living in the north were less likely to be cured. This result can be useful as a public health policy, since it is possible to prioritize this region to improve TB cure in MRJ. Possibly, the regions with the greatest probability of cure are associated with greater coverage of the family health strategy, such as in the western part of the municipality, covering more than 90% since 2010, in addition to Rocinha with 100% coverage since 2012. On the other hand, for the rest of the southern zone and the Tijuca region, it is probably due to much easier access to other health facilities, better socioeconomic situation and retention of physicians with good qualification.

From the spatial analysis, it was possible to perceive, as expected, an association between tuberculosis cure and better socioeconomic conditions such as educational level and income. Despite the association between race / color and cure in bivariate analysis, no association was found in the final spatial analysis model. Possibly, this variable has bad data entry problems. The spatial model demonstrated that the higher the educational level, the better the probability of cure, which is consistent with that expected from the literature [17, 19].

There was an apparent paradox between the lower probability of cure among the elderly, when analyzing the variable age at the individual level and the higher probability of cure the higher the aging rate at the level of the census tract. This may be due to the fact that, at the individual level, older patients have lower adherence to treatment due to intrinsic factors, leading to more adverse effects, increased drug interaction, forgetfulness of taking medication and less immunity. On the other hand, those patients residing in census tracts with higher rates of aging have a greater likelihood of cure, probably because this variable represents more structured communities from the socioeconomic viewpoint, which could have a longer life expectancy, and from a temporal viewpoint, because they are possibly older migration communities and with a larger social support network that would favor a better treatment and consequently a better probability of cure. It is worth noting that the aging rate was the variable that showed the highest probability of cure (OR 9.39), albeit with the greatest variability (95% CI 1.03-85.26), evidencing the importance of the social context for TB cure. The positive serology for HIV and alcohol abuse variables were associated with a lower probability of TB cure. In addition to the expected lower immunity, these patients may show less adherence, due to drug interaction and greater occurrence of medication side effects [37, 38].

The search for contactants was associated with a greater likelihood of cure for tuberculosis. Possibly, those cases whose professionals have made active search for contactants are the well-cared patients, receiving the greatest attention from the professionals.

It can be observed that the longer the implantation time of family health teams since the diagnosis of the disease, the greater the probability of TB cure, except for the category of 41 months or more, which evidenced the worst probability of cure among the categories. Data of this study are insufficient to explain this phenomenon, since it would be expected that teams with 41 months or more would be more likely to cure. One possible explanation is that the first family health teams in the MRJ were deployed in very vulnerable regions and, despite SMS-Rio efforts, some of them remained for a long time without being complete due to lack of doctors. This type of study does not allow a correlation between cause and effect and the external validity may not be reached. Other studies are necessary to confirm or not the results founded.

## Conclusion

The spatial analysis of cases of tuberculosis cure in the years 2012 to 2014 showed a significant spatial association with the cure of tuberculosis. In addition, regions in the municipality with more probability of cure were evidenced, which favors the manager to implement more efficient TB control measures. Regarding PHC coverage, a significant association was found between TB cure and time of family health implantation when the socioeconomic variables were controlled. This finding corroborates the importance of treating tuberculosis in this level of care.

## Supporting Information Legends

**S1 Checklist: STROBE Checklist**

**S2 Fig 3. Model of analysis for the cure of tuberculosis.** To cure tuberculosis, an individual with the disease must have been diagnosed and treatment performed. The environment, individual factors, access to health services and social condition (agglomeration, income, education, household conditions and demographic factors) of the individual influence the transmission, diagnosis and treatment of tuberculosis. Proper treatment results in the cure of tuberculosis.

**S3 Fig 4. Kernel ratio among new cases of tuberculosis and population. Municipality of Rio de Janeiro (RJ), Brazil, from 2012 to 2014.** Source: Cartographic data: IPP, Epidemiological data: SINAN-TB. Projection GCS. Datum WGS84

**S4 Fig 3. Map of probability of cure of tuberculosis using the Generalized Additive Model – GAM for cases (not cured) and controls (cure). Rio de Janeiro, 2012 to 2014.** Source: Cartographic data: IPP, Epidemiological data: SINAN-TB. Projection GCS. Datum WGS84

**S5 Fig 4. Final spatial analysis model using the generalized additive model - GAM for non-cure of tuberculosis. Rio de Janeiro, 2012 to 2014**

